# SARS-CoV-2 Omicron spike H655Y mutation is responsible for enhancement of the endosomal entry pathway and reduction of cell surface entry pathways

**DOI:** 10.1101/2022.03.21.485084

**Authors:** Mizuki Yamamoto, Keiko Tomita, Youko Hirayama, Jun-ichiro Inoue, Yasushi Kawaguchi, Jin Gohda

## Abstract

The SARS-CoV-2 Omicron variant reportedly displays decreased usage of the cell surface entry pathway mediated by the host transmembrane protease, serine 2 (TMPRSS2) and increased usage of the endosomal entry pathway mediated by cathepsin B/L. These differences result in different cell tropisms and low fusogenicity from other SARS-CoV-2 variants. Recent studies have revealed that host metalloproteases are also involved in cell surface entry and fusogenic activity of SARS-CoV-2, independent of TMPRSS2. However, the involvement of metalloproteinase-mediated cell entry and fusogenicity in Omicron infections has not been investigated. Here, we report that Omicron infection is less sensitive to the metalloproteinase inhibitor marimastat, like the TMPRSS2 inhibitor nafamostat, and is more sensitive to the cathepsin B/L inhibitor E-64d than infections with wild-type SARS-CoV-2 and other variants. The findings indicate that Omicron preferentially utilizes the endosomal pathway rather than cell surface pathways for entry. Moreover, the Omicron variant also displays poor syncytia formation mediated by metalloproteinases, even when the S cleavage status mediated by fusion-like proteases is unchanged. Intriguingly, the pseudovirus assay showed that a single mutation, H655Y, of the Omicron spike (S) is responsible for the preferential entry pathway usage without affecting the S cleavage status. These findings suggest that the Omicron variant has altered entry properties and fusogenicity, probably through the H655Y mutation in its S protein, leading to modulations of tissue and cell tropism, and reduced pathogenicity.

**Author summary:** Recent studies have suggested that the SARS-CoV-2 Omicron variant displays altered cell tropism and fusogenicity, in addition to immune escape. However, comprehensive analyses of the usage of viral entry pathways in Omicron variant have not been performed. Here, we used protease inhibitors to block each viral entry pathway mediated by the three host proteases (cathepsin B/L, TMPRSS2, and metalloproteinases) in various cell types. The results clearly indicated that Omicron exhibits enhanced cathepsin B/L-dependent endosome entry and reduced metalloproteinase-dependent and TMPRSS2-dependent cell surface entry. Furthermore, the H655Y mutation of Omicron S determines the relative usage of the three entry pathways without affecting S cleavage by the host furin-like proteases. Comparative data among SARS-CoV-2 variants, including Omicron, may clarify the biological and pathological phenotypes of Omicron but increase the understanding of disease progression in infections with other SARS-CoV-2 variants.

## Introduction

The Omicron variant of severe acute respiratory syndrome-associated coronavirus-2 (SARS-CoV-2), the causative virus of coronavirus disease 2019 (COVID-19), has spread globally since it was recognized as a variant of concern (VOC) by the World Health Organization on November 24, 2021. Despite its high infectivity, the pathogenicity of Omicron seems to be lower than that of other SARS-CoV-2 variants [1-4]. The Omicron variant has a large number of mutations in the viral genome, especially in the spike (S) region, which appears to contribute to the rapidly increasing number of cases worldwide by evading host immune responses in vaccinated or previously exposed individuals. Several research groups have reported that Omicron escapes from the neutralizing activities of therapeutic monoclonal antibodies and vaccine-elicited antibodies [5-9]. In addition to immune response escape, Omicron appears to exhibit different cell tropism from other SARS-CoV-2 variants [10-13], which may also contribute to the rapid spread of Omicron infection.

The S protein of SARS-CoV-2 is essential for viral entry into target cells. Prior to viral entry, the S protein is cleaved into S1 and S2 by host furin-like proteases before the release of viral particles from infected cells [14-16]. The first step is the binding of the S protein to the cell surface receptor angiotensin-converting enzyme 2 (ACE2) through the receptor binding domain (RBD) in the S1 domain [17, 18]. In the second step, the S2 domain is further cleaved into S2’ by either transmembrane serine protease 2 (TMPRSS2) localized in the plasma membrane or cathepsin B/L in endosomal vesicles. This step is called priming [19, 20]. S2’ priming causes membrane fusion between the viral particle and the plasma or endosomal membrane through interaction between the fusion peptide region and the cellular membrane, resulting in the release of viral RNA into the cytoplasm and its replication. We and others recently showed that host metalloproteinases are also involved in cell surface entry, independent of transmembrane protease, serine 2 (TMPRSS2) [21-23]. These results suggest that metalloproteinases act as the third host proteases for SARS-CoV-2 entry. Additionally, the S protein localized in the cell membrane is primed by TMPRSS2 or metalloproteinases after binding to ACE2 on the surface of neighboring cells, resulting in cell–cell fusion to generate syncytia. Syncytia formation is attributed to tissue damage [24-26].

A difference in the use of entry pathways has been observed among SARS-CoV-2 variants. The Delta variant, which was the predominant SARS-CoV-2 strain before the spread of Omicron, displays increased usage of the TMPRSS2-mediated cell surface entry pathway compared to wild-type (WT) SARS-CoV-2 [26]. In addition, the Delta variant shows enhanced syncytia formation, which may be related to its more severe pathogenicity compared to other variants [26, 27]. In contrast, the Omicron variant shows decreased usage of TMPRSS2-dependent cell surface entry [2, 10, 12, 13]. Furthermore, Omicron has low fusogenicity mediated by TMPRSS2, forming smaller syncytia than WT and Delta viruses [2, 10, 12, 13]. The reduction in TMPRSS2-mediated fusogenic activity may partially explain the low pathogenicity of the Omicron variant. However, the involvement of metalloproteinase-mediated viral entry and syncytia formation in Omicron infections has not been fully elucidated. In addition, the function of Omicron S mutations on entry pathway usage and fusogenicity remains unknown.

In this study, we investigated the relative usage of the three entry pathways— metalloproteinase-, TMPRSS2-, and cathepsin B/L-dependent pathway—by the Omicron variant. Our comprehensive analyses showed that Omicron preferentially utilized the cathepsin B/L-dependent endosomal entry pathway rather than the metalloproteinase- and TMPRSS2-dependent cell surface entry pathways in various cell types. Furthermore, syncytia formation mediated by metalloproteinases was significantly reduced, similar to that by TMPRSS2. Finally, H665Y of the Omicron S protein was identified as the mutation responsible for determining the preferential usage of the endosomal pathway without affecting S1/S2 cleavage status.

## Results

### Omicron variant preferentially uses the cathepsin B/L-dependent endosomal pathway over metalloproteinase- or TMPRSS2-dependent cell surface pathway

To evaluate the usage of the three different entry pathways by the SARS-CoV-2 variants, we used three protease inhibitors: nafamostat, marimastat, and E-64d. Nafamostat is a serine protease inhibitor that specifically blocks TMPRSS2-dependent (TMPRSS2 pathway) cell surface entry [28, 29]. Marimastat is a broad metalloproteinase inhibitor that blocks metalloproteinase-dependent (metalloproteinase pathway) cell surface entry [21]. E-64d is a cathepsin B/L inhibitor that suppresses the cathepsin B/L-dependent endosomal entry pathway (endosomal pathway) [19].

We first assessed viral replication of WT SARS-CoV-2 and an Omicron BA.1 variant in three cell types that have different entry pathways for SARS-CoV-2 infection. Intracellular viral RNA replication in Calu-3 airway epithelial cells after infection with WT (NCGM02) was severely blocked by nafamostat, but not by E-64d or marimastat (SFig. 1A). The findings indicated that WT infection predominantly depends on the TMPRSS2 pathway in Calu-3 cells. Infection of HEC50B endometrium cells was sensitive to E-64d and marimastat, but not to nafamostat (SFig. 1A). These findings indicated that WT infects HEC50B cells via both the endosomal and metalloproteinase pathways. Infection of OVISE ovary cells with WT was highly sensitive only to E-64d treatment, revealing that viral entry into OVISE cells predominantly depends on the endosomal pathway (SFig. 1A). The amount of intracellular viral RNA in the three cell lines was measured after 24 and 48 h of infection with WT or the Omicron variant (TY38-873) in Fig. 1A. Titers of WT and Omicron were measured using VeroE6-TMPRSS2 (JCRB) cells, which predominantly use the endosomal pathway, like OVISE cells (SFig. 1B). Compared to WT, Omicron replication was reduced in Calu-3 and HEC50B cells, which use TMPRSS2- or metalloproteinase-dependent cell surface pathways. In contrast, in OVISE cells, which predominantly utilize the endosomal pathway, WT and Omicron replicated similarly in accordance with the multiplicity of infection (MOI) measured using VeroE6-TMPRSS2 (JCRB) cells (Fig. 1A). These results support the hypothesis of a difference in the usage of viral entry pathways between the WT and Omicron variants.

**Fig 1.**
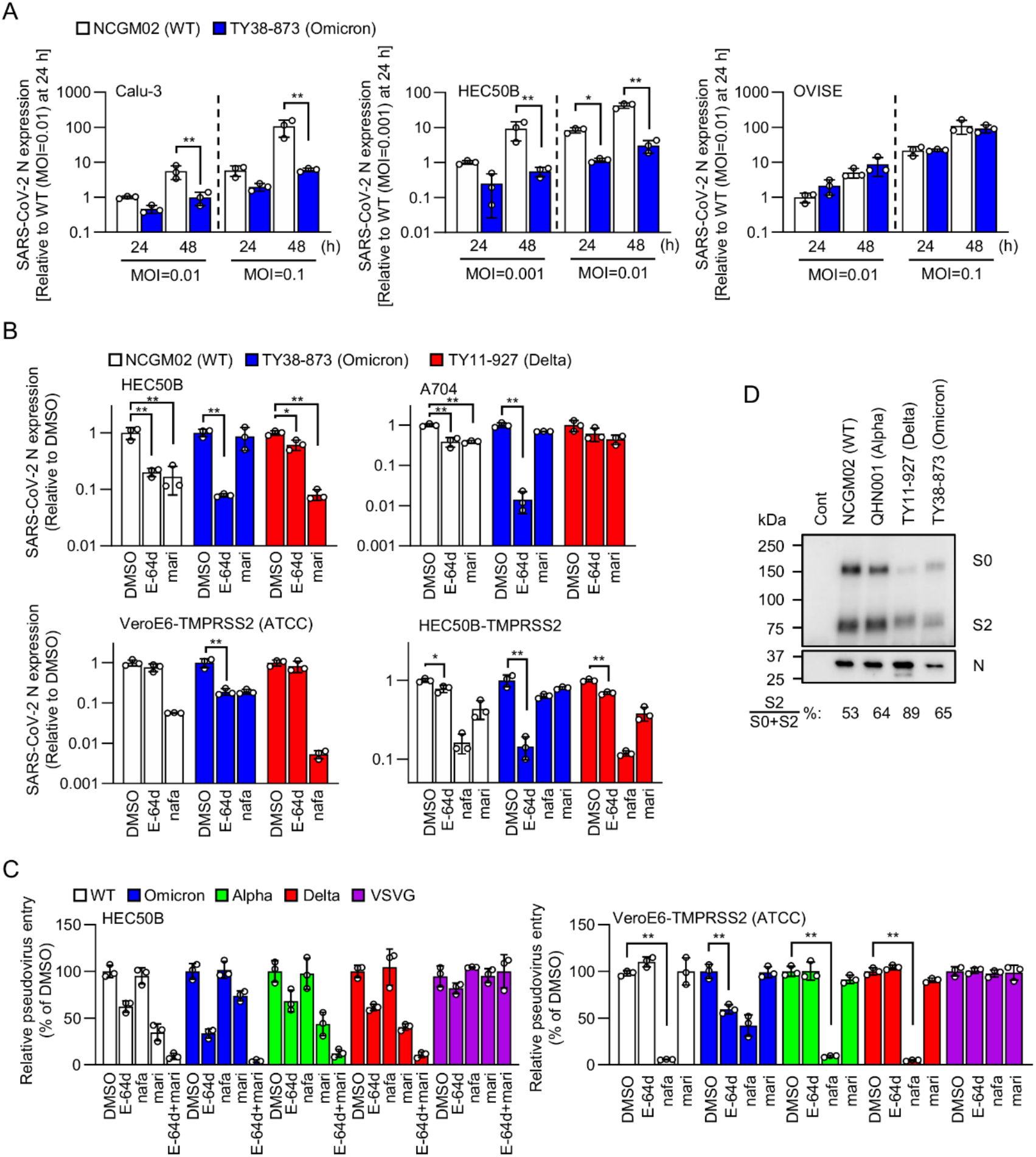
Omicron variant displays preferential usage of endosomal pathway over cell surface entry pathways. A. Comparison of replication rates between WT SARS-Cov-2 and Omicron variant in Calu-3, HEC50B, and OVISE cells. Intracellular viral RNA was measured by RT-qPCR after 24 h or 48 h infection of Calu-3, HEC50B, and OVISE cells with WT virus or Omicron variant at the indicated MOI. The relative amount of intracellular viral RNA was normalized to cellular *rpl13a* mRNA expression. Values are means ± SD (*n* = 3/group). B. Effects of protease inhibitors on infection with the SARS-CoV-2 variants. The cell lines were pretreated for 1 h with 25 μM E-64d, 10 μM nafamostat, or 1 μM marimastat. Intracellular viral RNA was measured by RT-qPCR after 24 h of infection of Calu-3 (MOI: 0.1), HEC50B (MOI: 0.01), A704 cells (MOI: 0.1), and HEC50B-TMPRSS2 cells (MOI: 0.01) with WT, Omicron, or Delta virus. The relative amount of intracellular viral RNA in each group was represented as ratio of the corresponding dimethyl sulfoxide (DMSO) control, which was set to 1, after being normalized to cellular *rpl13a* mRNA expression. C. Effects of protease inhibitors in HEC50B and VeroE6-TMPRSS2 (ATCC) on the entry of pseudoviruses bearing S proteins from SARS-CoV-2 WT or the variants. The relative pseudovirus entry was calculated by normalizing the luciferase activity for each condition to the luciferase activity of cells infected with pseudovirus in the presence of DMSO alone, which was set to 100%. Values are means ± SD (*n* = 3/group). *p > 0.05, **p > 0.01.

To compare the usage of the entry pathways by Omicron, WT, or Delta variant (TY11-927), we evaluated the effects of E-64d, nafamostat, and marimastat on infections in various cell types. In HEC50B cells, E-64d inhibited WT and Omicron infections, whereas Delta infection was less sensitive (Fig. 1B). In contrast, marimastat significantly suppressed WT and Delta infections, but not Omicron infections. In addition, in A704 human kidney cells, which use metalloproteinase and endosomal pathways for WT infection, similar to HEC50B cells (SFig. 1A), Omicron infection was much more dependent on the endosomal pathway than the WT and Delta infections (Fig. 1B). These data show that the metalloproteinase pathway is almost completely lost in Omicron infection, and that this variant significantly relies on the endosomal pathway. Next, we analyzed entry of Omicron into cells via the TMPRSS2 pathway. We tested VeroE6 cells ectopically expressing TMPRSS2 (VeroE6-TMPRSS2 (ATCC)), which predominantly depends on the TMPRSS2 pathway for WT infection (SFig. 1A). Although WT and Delta predominantly used the TMPRSS2 pathway in these cells, Omicron entry was significantly dependent on the endosomal pathway (Fig. 1B). We further evaluated the effects of protease inhibitors on the viral infection of HEC50B cells ectopically expressing TMPRSS2 (HEC50B-TMPRSS2), which have all three entry pathways for WT infection (Fig. 1B). In Omicron infection, the contribution of the endosomal pathway was enhanced, while the TMPRSS2 and metalloproteinase pathways were less preferentially utilized compared to WT and Delta (Fig. 1B). These findings suggest that Omicron preferentially utilizes the endosomal pathway, even when the target cells have all the three pathways. In addition to the live virus experiment, we used replication-incompetent vesicular stomatitis virus (VSV) pseudoviruses bearing the S protein derived from WT, Omicron, Alpha, or Delta. Consistent with the live virus experiments, the Omicron S pseudovirus exhibited increased usage of the endosomal pathway and decreased usage of the metalloproteinase pathway in HEC50B and A704 cells compared to WT, Alpha, and Delta S pseudoviruses (Fig. 1C and SFig. 2). In addition, increased usage of the endosomal pathway by the Omicron S pseudovirus was observed in both VeroE6-TMPRSS2 (ATCC) cells (Fig 1C) and Caco-2 (colon cells) harboring the TMPRSS2 and endosomal pathways (SFig. 2). Taken together, these results suggest that Omicron S preferentially uses the cathepsin B/L-dependent endosomal entry pathway, rather than metalloproteinase- and TMPRSS2-dependent cell surface entry pathways. Interestingly, there was no difference in the dose-dependency of the inhibitory effects of nafamostat between Omicron and the other variants in Calu-3 cells (SFig.1 C and SFig. 2). In OVISE cells, WT and Omicron infections showed similar sensitivities to E-64d (SFig. 2). These data suggest that Omicron uses the same entry pathway as WT when infecting cells that have only one entry pathway.

**Fig 2.**
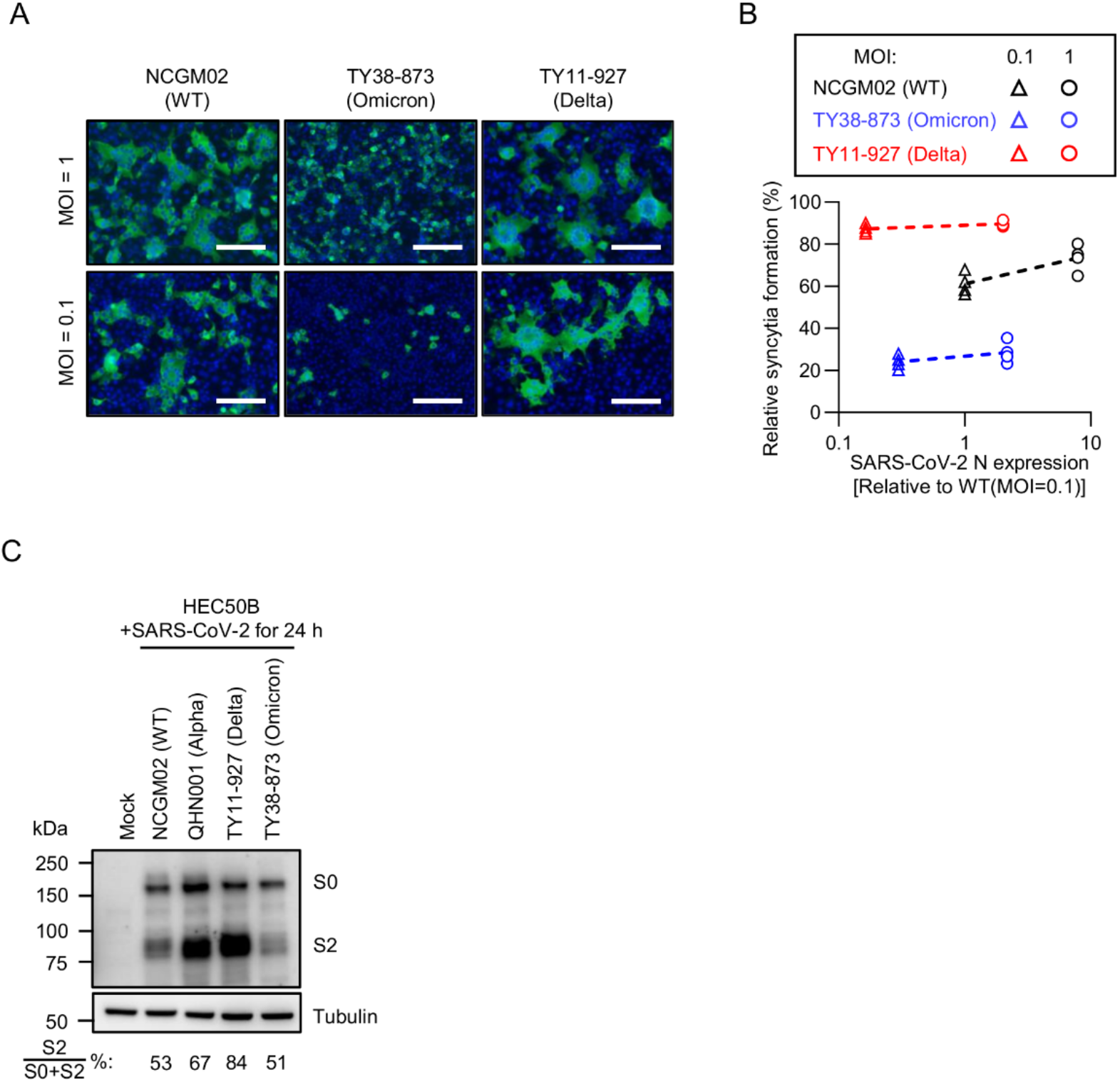
Omicron variant displays poor syncytium formation mediated by metalloproteinases. A. Syncytia formation of HEC50B cells induced by infection with SARS-CoV-2 (NCGM02, TY11-927 or TY38-873). Cells were infected with SARS-CoV-2 at an MOI of 0.1 or 1. After 24 h infection, cells were fixed and stained with anti-SARS-CoV-2 N antibody (green). The cell nuclei were stained with Hoechst 33342 (blue). White bar = 200 mm. B. Quantification of syncytia shown in A and amount of viral RNA in infected HEC50B cells. C. Expression of S protein in HEC50B cells. Cells were infected with SARS-CoV-2 (NCGM02, QHN001, TY11-927 or TY38-873) for 24 h and cells lysed for western blotting analysis. S0: uncleaved S protein; S2: cleaved S2 domain of the S protein.

Cleavage of SARS-CoV-2 S protein by host furin-like proteases affects the entry pathways for SARS-CoV-2 infection [2, 12, 13, 26, 30]. Therefore, we investigated the cleavage status of the S proteins in the variants. Western blot analysis of the virions using an anti-S2 domain antibody showed that the S protein of the Alpha variant was cleaved to the same degree as the WT S, while the cleavage efficiency of the Delta S was higher than that of the WT and Alpha S (Fig. 1D). Interestingly, there was no significant difference in the cleavage efficiency between Omicron S and WT S (Fig. 1D). These results suggest that Omicron S can change the relative usage of entry pathways, even when Omicron S is cleaved as efficiently as WT S.

### Omicron variant displays poor syncytium formation mediated by metalloproteinases without affecting the S1/S2 cleavage of its S protein

Reduced syncytia formation through the TMPRSS2 pathway has been reported in Omicron, suggesting a correlation with lower pathogenicity [2, 10, 12, 13]. We previously reported that metalloproteinases can also mediate syncytia formation by WT virus in HEC50B cells [21]. Thus, we examined the ability of Omicron to form syncytia via metalloproteinases in these cells. After 24 h of infection with WT, Omicron, or Delta, the cells were stained with anti-SARS-CoV-2 N protein to observe the formation of syncytia. WT infection formed numerous syncytia both at a low MOI of 0.1 and a high MOI of 1 (Fig. 2A and SFig. 3). Consistent with our previous report [21], metalloproteinase-mediated syncytia formation was observed in WT infection at both MOI values (SFig. 3). However, Omicron infection induced much smaller and fewer syncytia than WT infection, even when the cells were infected at the high MOI of 1 (Fig. 2A). Although Delta infection also induced syncytia formation in a metalloproteinase-dependent manner in HEC50B cells (SFig. 3), the syncytia that formed where larger than those in WT infection (Fig. 2A), as enhanced syncytia formation in TMPRSS2 expressing cells reported previously [26, 27]. As Omicron replication was less than that of WT virus in HEC50B cells (Fig. 1A), it is possible that poor syncytia formation by Omicron was due to its reduced replication in HEC50B cells. Therefore, we measured the intracellular viral RNA levels when syncytia formation was observed. Omicron infection at an MOI of 1 resulted in much poorer syncytia formation than WT infection at an MOI of 0.1, even when the Omicron infection produced more viral RNA in the cells than the WT infection (Fig. 2B). In contrast, Delta infection produced more syncytia than WT and Omicron infections, regardless of the amount of viral RNA (Fig. 2B). These results indicated that the Omicron variant has a poor ability to form syncytia by TMPRSS2 and metalloproteinases, and that the Delta variant induces enhanced syncytia formation by metalloproteinases.

**Fig 3.**
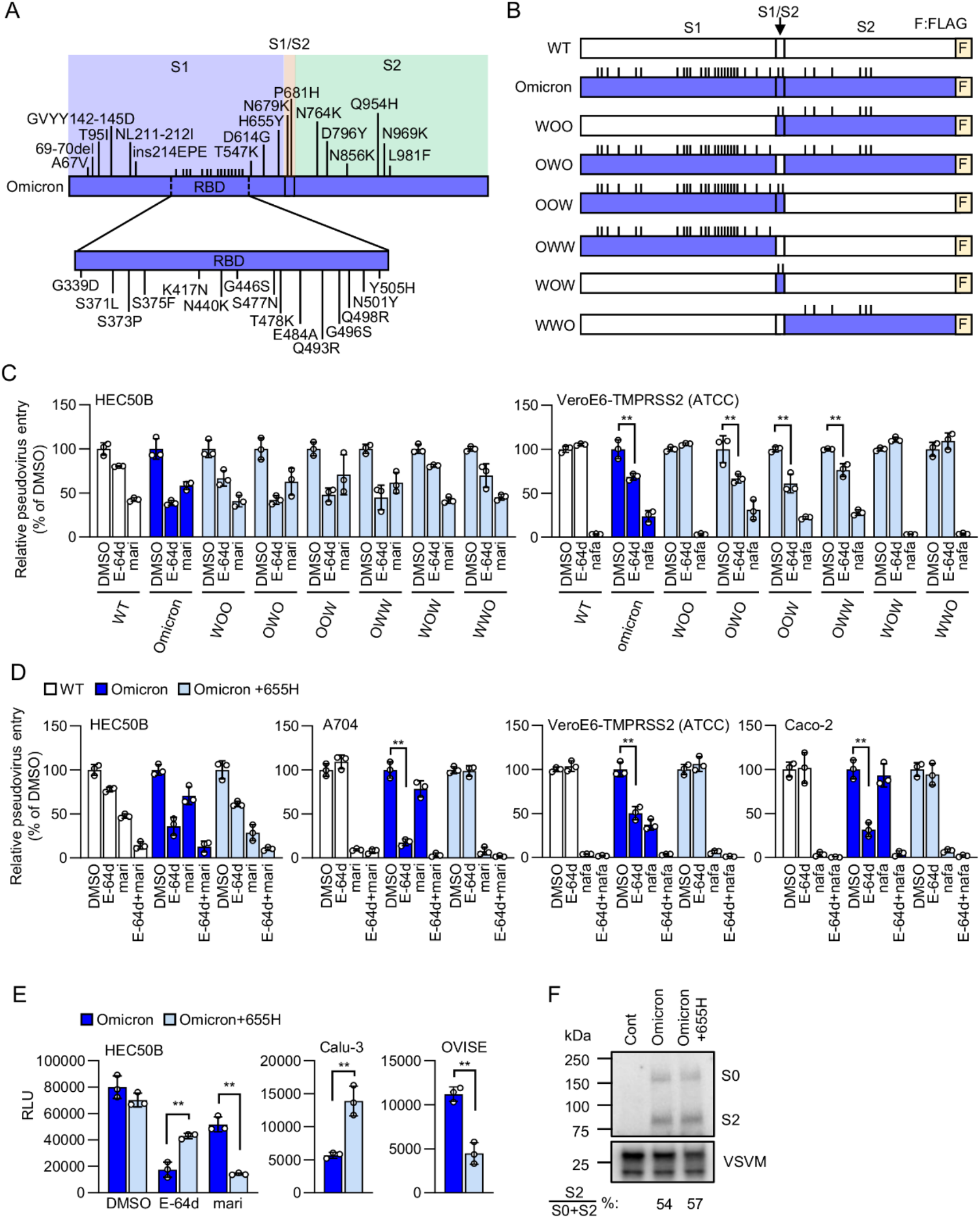
H655Y mutation in Omicron S protein facilitates viral entry through the endosomal pathway. A. Schematic illustration of mutations in Omicron S protein. S1 (Met^1^ to Thr^676^), S1/S2 boundary (Gln^677^ to Arg^685^), and S2 domain (Ser^686^ to Thr^1273^). Numbers refer to the amino acid residues corresponding to WT S protein. B. Schematic illustration of C-terminally Flag-tagged chimeric S proteins in which the S1, S1/S2 boundary, and S2 domain from the WT S (white) or the Omicron S (blue) are indicated. F: Flag-tag. C. Effects of protease inhibitors in HEC50B and VeroE6-TMPRSS2 (ATCC) cells on the entry of pseudoviruses bearing chimeric S proteins. D. Effects of protease inhibitors in HEC50B, A704, VeroE6-TMPRSS2 (ATCC) and Caco-2 cells on the entry of pseudoviruses bearing Omicron S or Omicron S protein with tyrosine residue at position 655 substituted to histidine (Omicron +665H). E. Infectivity of pseudoviruses bearing Omicron S or Omicron S+665H in HEC50B, Calu3 and OVISE cells. F. Expression of chimeric S protein in pseudoviruses. S proteins were detected using an anti-Flag-tag antibody that binds to a Flag-tag on the C-terminus of the S proteins (top). Detection of the vesicular stomatitis virus matrix protein (VSV M) served as the control (bottom). S0: uncleaved S protein; S2: cleaved S2 domain of the S protein. The relative pseudovirus entry was calculated by normalizing the luciferase activity for each condition to the luciferase activity of cells infected with pseudovirus in the presence of DMSO alone, which was set to 100% in C, D and E. Values are means ± SD (*n* = 3/group). **p > 0.01.

Several studies reported that S1/S2 cleavage of the SARS-CoV-2 S protein expressed in infected cells affects syncytium formation [2, 12, 13]. However, in the present study, there was no significant difference in the S1/S2 cleavage status between the WT, Alpha, and Omicron variants in HEC50B cells, although Delta S was cleaved more than the S proteins of the other variants (Fig. 2C), as previously reported [26]. These results indicate that Omicron forms syncytia poorly, even when S cleavage status is similar to that of WT.

### H655Y mutation of Omicron spike protein determines preferential usage of endosomal pathway without affecting S1/S2 cleavage status

To elucidate the molecular mechanism underlying the preferential usage of the endosomal pathway by Omicron, we attempted to identify the responsible mutation sites in the S protein. The Omicron BA.1 variant (TY38-873) has 32 amino acid mutations in its S protein compared to the WT, with 24 mutations in the S1 region, including the RBD, two mutations in the S1/S2 boundary region including the furin cleavage site, and six mutations in the S2 region (Fig. 3A). We generated various pseudoviruses with chimeric S proteins that consisted of the S1, S1/S2 boundary, and S2 regions derived from either WT or Omicron (Fig. 3B). In HEC50B cells, pseudoviruses bearing the S proteins consisting of the WT S1 region (WOO, WOW, and WWO) infected the cells mainly through the metalloproteinase pathway (Fig. 3C), similar to the WT pseudovirus. In contrast, the pseudoviruses bearing the S proteins consisting of the Omicron S1 region (OWO, OOW, and OWW) infected the cells mainly through the endosomal pathway (Fig. 3C). Furthermore, in VeroE6-TMPRSS2 (ATCC) cells, the WOO, WOW, and WWO pseudoviruses predominantly used the TMPRSS2 pathway, similar to the WT pseudovirus, whereas the OWO, OOW, and OWW pseudoviruses showed the same pattern of entry pathway usage as the Omicron S pseudovirus (Fig. 3C). These results clearly demonstrated that the S1 region of the Omicron S protein is responsible for preferential usage of the endosomal pathway.

To identify the amino acid mutations in the S protein responsible for the preferential usage of the endosomal pathway by Omicron, we generated 24 pseudoviruses bearing Omicron S proteins, with each mutated amino acid residue in the Omicron S1 region substituted for the corresponding amino acid in the WT S1 region. Among them, the Omicron S protein with an amino acid substitution at position 373 (373S) failed to produce infectious virus particles. Therefore, 23 pseudoviruses were used for subsequent analyses. Only the pseudovirus bearing the Omicron S protein with a tyrosine residue at position 655 substituted for histidine (Omicron S +665H) out of the 23 pseudovirus infected HEC50B cells predominantly through the metalloproteinase pathway as in the WT pseudovirus (SFig. 4). In VeroE6-TMPRSS2 (ATCC) cells, only the (Omicron S +655H) pseudovirus predominantly used the TMPRSS2 pathway, similar to the WT pseudovirus (SFig. 4). In addition, the Omicron S +655H pseudovirus infected A704 and Caco-2 cells predominantly through the cell surface pathways, as well as HEC50B and VeroE6-TMPRSS2 (ATCC) cells (Fig. 3D). These results indicate that tyrosine at position 655 of the Omicron S protein is responsible for preferential usage of the cathepsin B/L-dependent endosomal entry pathway over the metalloproteinase- or TMPRSS2-dependent cell surface entry pathway.

To investigate whether the substitution of tyrosine for histidine at position 655 in the Omicron S protein enhances the cell surface pathways or impairs the endosome pathway, we analyzed the effect of the 655H substitution on infectivity through each pathway. Luciferase activities from HEC50B cells infected with Omicron and the (Omicron S +655H) pseudoviruses were almost the same (Fig. 3E). The Omicron S +655H pseudovirus infected cells more efficiently than the Omicron pseudovirus when the cathepsin B/L-dependent endosome pathway was blocked by E-64d (Fig. 3E). In contrast, the Omicron +655H pseudovirus was less infectious than the Omicron pseudovirus when the metalloproteinase pathway was blocked by marimastat (Fig. 3E). Moreover, the Omicron S +655H pseudovirus was more infectious to Calu-3 cells, which have only the TMPRSS2 pathway, and less infectious to OVISE cells, which have only the endosome pathway, than was the Omicron pseudovirus. Western blot analysis of the S proteins on the pseudoviruses revealed no significant difference in the cleavage status between Omicron S and Omicron S +655H (Fig. 3F). These results strongly suggest that the H655Y mutation of Omicron S is responsible for alteration of the usage of the entry pathways by facilitating entry through the cathepsin B/L-dependent endosome pathway and impairing entry through the metalloproteinase- and TMPRSS2-dependent cell surface pathways without affecting the S1/S2 cleavage status.

## Discussion

We recently reported that host metalloproteinases are involved in cell surface fusion for SARS-CoV-2 entry, independent of TMPRSS2, and that the pathway SARS-CoV-2 uses for cell surface entry is cell type-dependent [21]. Although tissues and cells preferentially use the metalloproteinase pathway for SARS-CoV-2 infection *in vivo* are still unknown, reduced usage of the metalloproteinase pathway by Omicron could contribute to the modulation of tissue and cell tropism. Moreover, our data clearly show that Omicron displays a significantly reduced metalloproteinase-dependent fusogenic activity. Given that the fusogenicity of SARS-CoV-2 correlates well with disease severity, the low fusogenic activity of Omicron for metalloproteinase dependence is likely to contribute to the less severe disease phenotype of Omicron in cooperation with the low fusogenicity mediated by TMPRSS2. Conversely, the high fusogenic activity mediated by metalloproteinases, as observed in Delta (Fig. 2AB), may lead to severe pathogenicity of SARS-CoV-2. Thus, fusogenicity mediated by metalloproteinases, as well as TEMPRSS2, may be useful as a barometer for the pathological severity of SARS-CoV-2 variants.

In this study, we successfully identified a single mutation, H655Y, in the Omicron S protein, which is responsible for the altered entry properties. Furthermore, our data reveal that the H655Y mutation facilitates entry through the endosomal pathway and impaired entry through the cell surface pathways (Fig. 3E). H655Y is a common mutation that is present in BA.1, BA.2, and BA.3, variants classified as Omicron, and is considered to be an important mutation that defines the characteristics of Omicron. in the WT virus, the H655Y mutation has been reported in a variety of SARS-CoV-2 infection studies using cell culture and animal models [31-35]. Despite these facts, only a few studies have explored the impact of H655Y mutation on biological and pathological phenotypes. One research group reported that the H655Y mutation in the WT virus enhances viral replication and transmission, possibly through increased S1/S2 cleavage [35]. According to these results, the H655Y mutation in the WT S backbone appears to have different effects on viral phenotypes than that in the Omicron S backbone. In addition, the H665Y mutation was observed in the gamma (P.1) variant among VOCs. However, the role of this mutation in the phenotype of the gamma variant has not yet been investigated. Further studies are needed to assess how the H655Y mutation affects the biological and pathological phenotypes of WT virus and other VOCs than the Omicron.

Inhibition of S1/S2 cleavage has been reported to reduce the usage of the TMPRSS2 and metalloproteinase pathways and increases the usage of the endosomal pathway [15, 21, 30, 36]. Furthermore, recent studies reported that the Omicron S protein expressed in some cell types, such as VeroE6-TMPRSS2 and HEK293 cells, is poorly cleaved into S1 and S2 compared to the WT S, resulting in a reduction in the fusogenic activity of Omicron [2, 12, 13]. However, we did not observe any significant differences in the cleavage of the S protein in the virions and infected HEC50B cells between Omicron S and WT S cells, even when the cell surface entry pathway and fusogenic activity were markedly suppressed (Fig. 1D and 2C). These data strongly suggest that Omicron S has the ability to change the entry pathway usage and fusogenicity, regardless of the S1/S2 cleavage status, in some cell types used in this study. Importantly, pseudovirus analysis shows that the H655Y mutation of the Omicron S causes a change in entry pathway usage without affecting the S1/S2 cleavage status, strongly suggesting that H655Y regulates entry properties through mechanisms other than the reduction of the S1/S2 cleavage efficiency.

A putative mechanism is that the H655Y mutation alters the Omicron entry pathway by affecting the stability of the S1/S2 complex formed after cleavage by furin-like proteases. The H655Y mutation is localized in subdomain 2 (SD2) of the S1 domain. Structural studies of the SARS-CoV-2 S protein indicate that SD2 is proximal to the S2 domain [37-39], suggesting that the stability of the S1/S2 complex may be influenced by some amino acid mutations in SD2. In fact, the D614G mutation in SD2 destabilizes the interface between S1 and S2, facilitating disorder of a region including the S2’ cleavage site, leading to increased infectivity [37, 39]. In contrast, the H655Y mutation in SD2 may make the S1/S2 complex more stable in the backbone of the Omicron S. According to the recently reported three-dimensional structure of Omicron S [40], the S trimer has a structure that is broadly common to that of WT S (SFig. 5AB). The C-terminal peptide of the S1 domain containing the H655Y mutation interacts with the N-terminal peptide of the S2 domain. Comparing the structures around amino acid position 655 in the spike protein of Omicron and WT, both spikes have hydrogen bonds between the two main chains (position 655-656 in S1 and position 694-695 in S2). Interestingly, Omicron S has an additional hydrogen bond between the side chains of the tyrosine residue at position 655 (Y655) in S1 and the threonine residue at position 696 (T696) in S2) (SFig. 5C). This additional hydrogen bond in Omicron S suggests that it may stabilize the S1/S2 complex. The increased stability possibly prevents TMPRSS2- and metalloproteinase-mediated S2’ priming on the cell surface by interrupting the exposure of the S2’ priming site. However, increased stability may have a positive effect on the endosomal entry of Omicron. For instance, a stable S1/S2 complex may increase the chance of endocytosis of viral particles by persistent anchoring of viral particles to the target cell membrane. In other words, stabilization of the binding between the S1 domain bound to ACE2 and the S2 domain anchored to the viral membrane may allow for efficient internalization of viral particles during endocytosis, which involves a dynamic change in cell membrane curvature. Elucidation of the differences between the Omicron S and the mutated S (Omicron +655H) by structural and biochemical analyses is needed to explain how the S H665Y mutation changes the relative usage of entry pathways in the Omicron variant.

The finding that the H655Y mutation of the Omicron S protein can change the usage of viral entry pathways might contribute to an understanding of the physiological and pathological phenotypes of the Omicron variant, and to the prediction of the pathogenesis of the next emerging variants from Omicron. Although the link between the altered usage of entry pathways and the rapid spread rate of Omicron is unknown, it is possible that if Omicron acquires mutations again around amino acid position 655 in the S protein, it may be a mutant strain that preferentially utilizes cell surface entry pathways, similar to WT and Delta, and could be seriously pathogenic with high infectivity. It may be necessary to monitor the occurrence of a mutation at tyrosine 655 in the S protein of Omicron during the current COVID-19 pandemic.

In summary, the findings of this study show that the Omicron variant preferentially uses the endosome pathway over cell surface pathways through the H655Y mutation in the S protein, even in cases where S cleavage by furin-like proteases is not affected. These results may contribute to the understanding of the biological and pathological features of the Omicron variant and the next emerging SARS-CoV-2 variants, and to the development of therapeutic approaches for COVID-19.

## Materials and Methods

### Cells and viruses

VeroE6 (CRL-1586), 293T (CRL-3216), A704 (HTB-45) and Calu-3 (HTB-55) cells were obtained from American Type Culture Collection (Rockville, MD, USA). OVISE (JCRB1043), HEC50B (JCRB1145), and VeroE6-TMPRSS2 (JCRB 1819) cells were obtained from the Japanese Collection of Research Bioresources Cell Bank (Osaka, Japan). Caco-2 cells (RCB0988) were obtained from RIKEN BioResource Research Center (Tsukuba, Japan). A704, Calu-3, VeroE6 (CRL-1586), HEC50B, and Caco-2 cells were maintained in Eagle’s minimum essential medium (EMEM; 055-08975, FUJIFILM Wako Pure Chemical, Osaka, Japan) containing 15% fetal bovine serum (FBS). VeroE6-TMPRSS2 (JCRB 1819) cells were cultured in DMEM containing 10% FBS and 1 mg/mL G418. OVISE cells were maintained in Roswell Park Memorial Institute (RPMI)-1640 medium (189-02025, FUJIFILM Wako Pure Chemical) containing 10% FBS. The WT SARS-CoV-2 isolate (UT-NCGM02/Human/2020/Tokyo)[41] was obtained from Dr. Yoshihiro Kawaoka, The University of Tokyo. An alpha isolate (B.1.1.7 linage, QHN001), Delta isolate (B.1.617.2 linage, TY11-927), and Omicron isolate (BA.1 linage, TY38-873) were obtained from the National Institute of Infectious Diseases, Japan. The viruses were propagated in VeroE6-TMPRSS2 (JCRB 1819) cells in DMEM containing 5% heat-inactivated FBS at 37 °C in 5% CO_2_. Briefly, SARS-CoV-2 was added to VeroE6-TMPRSS2 (JCRB 1819) cells and incubated for 30 min at 37 °C. The culture medium was then replaced with fresh medium. Before the supernatants were harvested, the cells were incubated for an additional 48 h (for UT-NCGM02, QHN001, and TY11-927) or 72 h (for TY38-873). The virus titer was determined by a plaque assay using VeroE6-TMPRSS2 (JCRB 1819) cells.

### Expression vector construction

Synthetic DNA corresponding to the codon-optimized S gene of SARS-CoV-2 (Wuhan-Hu-1, RefSeq: NC_045512.2), SARS-CoV-2 Omicron variants (BA.1, GISAID: EPI_ISL_7418017), and chimeric S gene (S1, S1/S2 boundary, and S2 domains were derived from either Wuhan-Hu-1 or Omicron variants) with DNA sequences corresponding to the Flag-tag 5-GGA GGC <u>GAT TAC AAG GAT GAC GAT GAC AAG</u> TAA-3’ (underline, Flag-tag) at the 3 end were generated by Integrated DNA Technologies (Coralville, IA, USA). Mutant Omicron S genes were constructed by PCR-based site-directed mutagenesis. To construct expression vectors for the S protein, the coding regions were cloned into a lentiviral transfer plasmid (CD500B-1; SBI, Palo Alto, CA, USA).

### Preparation of pseudotype VSV viral particles and infection experiments

VSV particles were prepared as previously described [21, 42]. For the infection assay, the target cells were seeded in 96-well plates (2 × 10^4^ cells/well) and incubated overnight. The cells were pretreated with inhibitors for 1 h before infection. Pseudotype viral particles were added to the cells in the presence of inhibitors. Luciferase activity was measured 16-h post-infection using the Bright-Glo Luciferase Assay System and a Centro xS960 luminometer (Berthold, Bad Wildbad, Germany).

### Quantification of intracellular SARS-CoV-2 RNA

Intracellular SARS-CoV-2 RNA was quantified as previously described [21]. Cell lysis and cDNA synthesis were performed using the SuperPrep II Cell Lysis & RT Kit (SCQ-401, TOYOBO, Osaka, Japan) following the manufacturer’s instructions. Viral RNA was detected using THUNDERBIRD SYBR qPCR Mix (TOYOBO) at 95 °C for 3 min, followed by 50 cycles of 95 °C for 10 s and 60 °C for 1 min. Fluorescence was detected using a CFX Connect^™^ Real-Time PCR Detection System (Bio-Rad, Hercules, CA, USA). The mRNA expression level of ribosomal protein L13a (Rpl13a) in each sample was used to standardize the data. The primer sequences used were 5’-AAATTTTGGGGACCAGGAAC-3’ as forward primer and 5’- TGGCAGCTGTGTAGGTCAAC-3’ as reverse primer for the SARS-CoV-2 N gene; 5’- TGTTTGACGGCATCCCAC-3’ as forward primer and 5’-CTGTCACTGCCTGGTACTTC- 3’as reverse primer for the human *Rpl13a* gene; and 5’-CTCAAGGTTGTGCGTCTGAA-3’ as forward primer and 5’-CTGTCACTGCCTGGTACTTCCA-3’ as reverse primer for the African green monkey *Rpl13a* gene.

### Western blotting

Western blotting was performed as described previously [21, 28]. The primary antibodies used were mouse anti-SARS-CoV-2 S2 domain (1:1000, GTX632604, GeneTex, Irvine, CA, USA), rabbit anti-Flag-tag (1:1000, PM020, MBL, Woburn, MA, USA), mouse anti-tubulin (1:1000, CP06, Millipore, Billerica, MA, USA), and mouse anti-VSVM (1:1000, 23H12, absolute antibody, Oxford, UK). The secondary antibodies used were horseradish peroxidase (HRP)-linked donkey anti-rabbit IgG (NA934; GE Healthcare) and HRP-linked donkey anti-mouse IgG (NA931V; GE Healthcare). Cell supernatants containing the viral particles were lysed for western blot analysis.

### Immunofluorescence staining

Immunofluorescence analysis of HEC50B cells was performed as described previously [21]. After 24 h infection, fixed cells were incubated with anti-SARS-CoV-2 nucleocapsid (1:1000, GTX135357) primary antibody for 16 h at 4 °C and detected with anti-rabbit-Alexa488 (1:200, A11008, Invitrogen, Carlsbad, CA, USA) secondary antibodies for 40 min at room temperature. Cell nuclei were stained with 1 μg/mL Hoechst 33342 (#080-09981, FUJIFILM Wako Pure Chemical). Fluorescent signals were detected using a BZ-X810 fluorescence microscope (Keyence, Osaka, Japan).

### Statistical analysis

Statistically significant differences between mean values were determined using a two-tailed Student’s t-test. Dunnett’s and Tukey’s tests were used for multiple comparisons. All data represent three independent experiments, and values represent the mean ± standard deviation, with p<0.05, considered statistically significant.

## Acknowledgments

We thank Yoshihiro Kawaoka for providing the SARS-CoV-2 isolates (UT-NCGM/Human/2020/Tokyo). This research was supported in part by the Japan Agency for Medical Research and Development (AMED) under Grant Number JP20wm0125002 to M.Y., Y.K., and J.G., by grants-in-aid from the Japanese Society for the Promotion of Science (18K15235 to M.Y.), AMED [Program of Japan Initiative for Global Research Network on Infectious Diseases (JGRID) JP20wm0125002 to Y.K.], and [Research on Development of New Drugs 21ak0101165 to M.Y.], and from the University of Tokyo (Promoting practical use of measures against coronavirus disease 2019 [COVID-19] to J.I.).

## Author Contributions

M.Y. and J.G. designed the study and prepared the manuscript. M.Y., K.T., Y.H., and J.G. performed experiments. M.Y., J.I., Y.K., and J.G. analyzed the data.

## Supporting information

**SFig 1.**
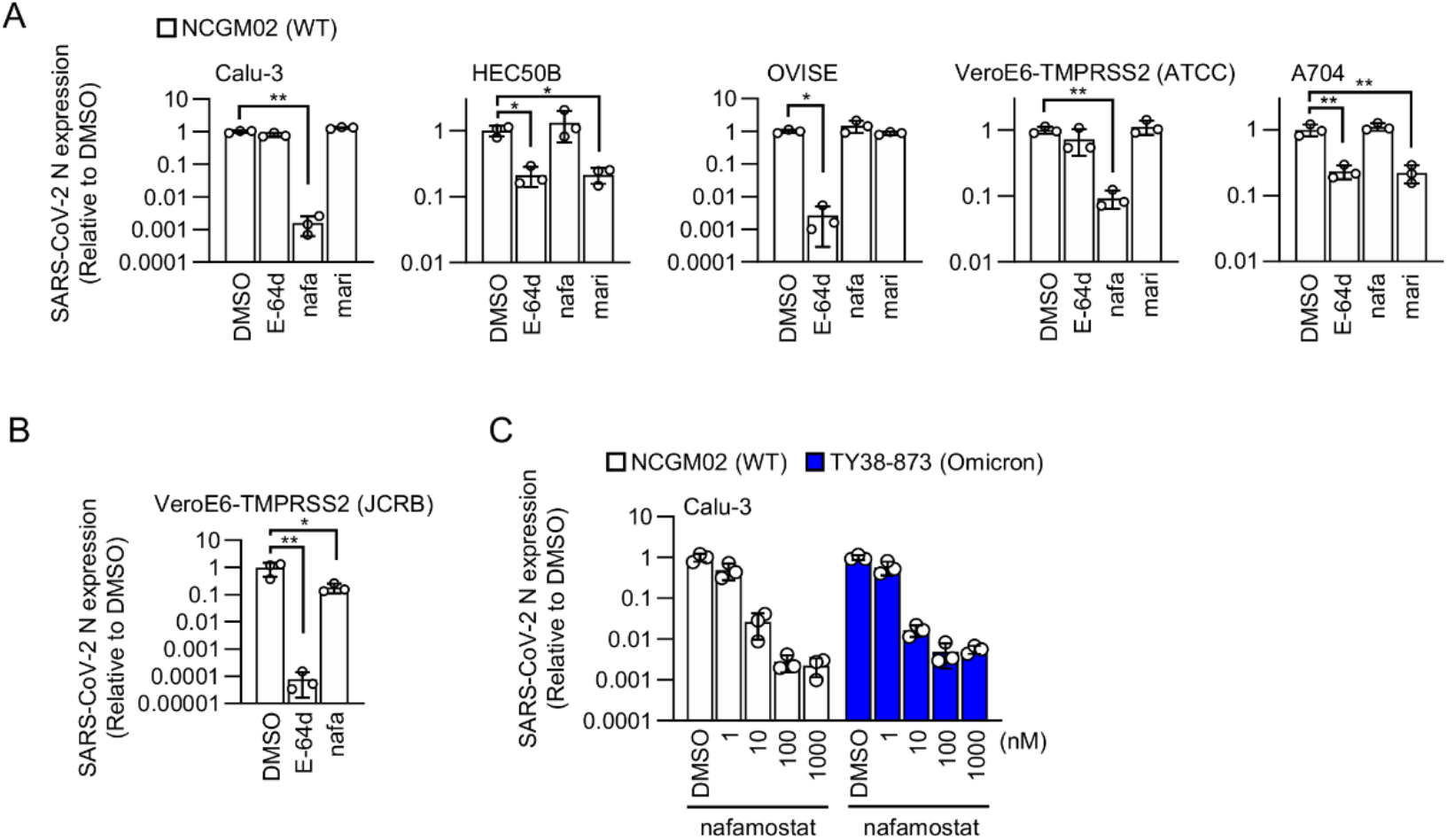
Effects of protease inhibitors on SARS-CoV-2 infection. A. The indicated cell lines were pretreated for 1 h with 25 μM E-64d, 10 μM nafamostat, or 1 μM marimastat. Intracellular viral RNA was measured by RT-qPCR 24 h after infection with Calu-3 (MOI: 0.1), HEC50B (MOI: 0.01), OVISE cells (MOI: 0.1), VeroE6-TMPRSS2 cells (MOI: 0.01), and A704 cells (MOI: 0.1) with WT virus. B. VeroE6-TMPRSS2 (JCRB) cells were pretreated for 1 h with 25 μM E-64d or 10 μM nafamostat. Intracellular viral RNA was measured by RT-qPCR 24 h after infection with WT virus. The relative amount of intracellular viral RNA in each group was represented as the ratio of the corresponding DMSO control, which was set to 1 after normalization to cellular *rpl13a* mRNA expression. C. Calu-3 cells were pretreated for 1h with nafamostat at the indicated doses. Intracellular viral RNA was measured by RT-qPCR after 24 h of infection with WT or Omicron at an MOI of 0.1. Values are presented as mean ± SD (*n* = 3/group). *p > 0.05, **p > 0.01.

**SFig 2.**
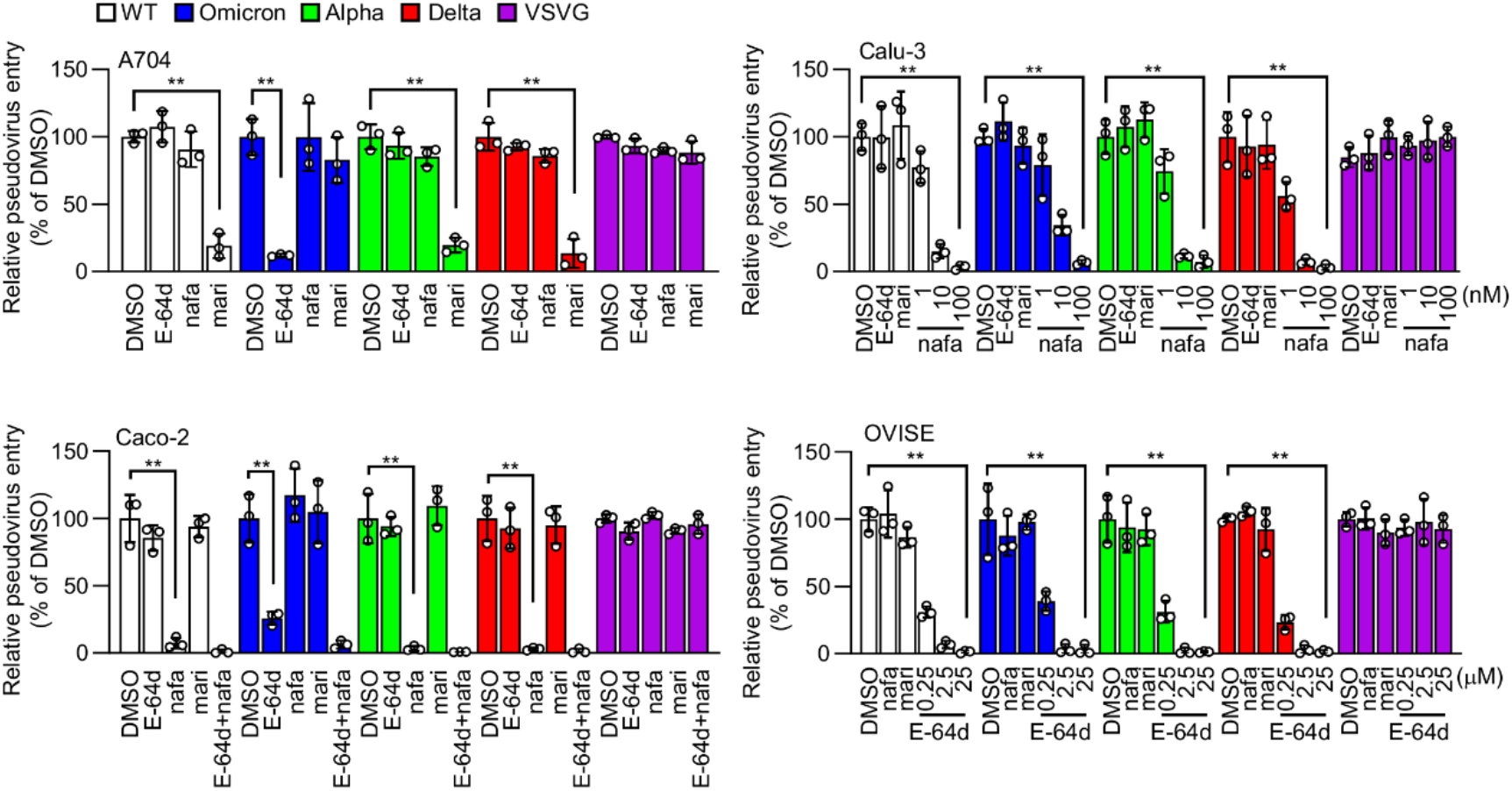
Effects of protease inhibitors on infection with VSV pseudoviruses bearing S proteins derived from SARS-CoV-2 variants. Effects of protease inhibitors in Calu-3, A704, OVISE, and Caco-2 cells on the entry of pseudoviruses bearing S proteins from SARS-CoV-2 WT or variants, or VSVG protein. The relative pseudovirus entry was calculated by normalizing the luciferase activity for each condition to the luciferase activity of cells infected with pseudovirus in the presence of DMSO alone, which was set to 100%. Values are presented as mean ± SD (*n* = 3/group). *p > 0.05, **p > 0.01.

**SFig 3.**
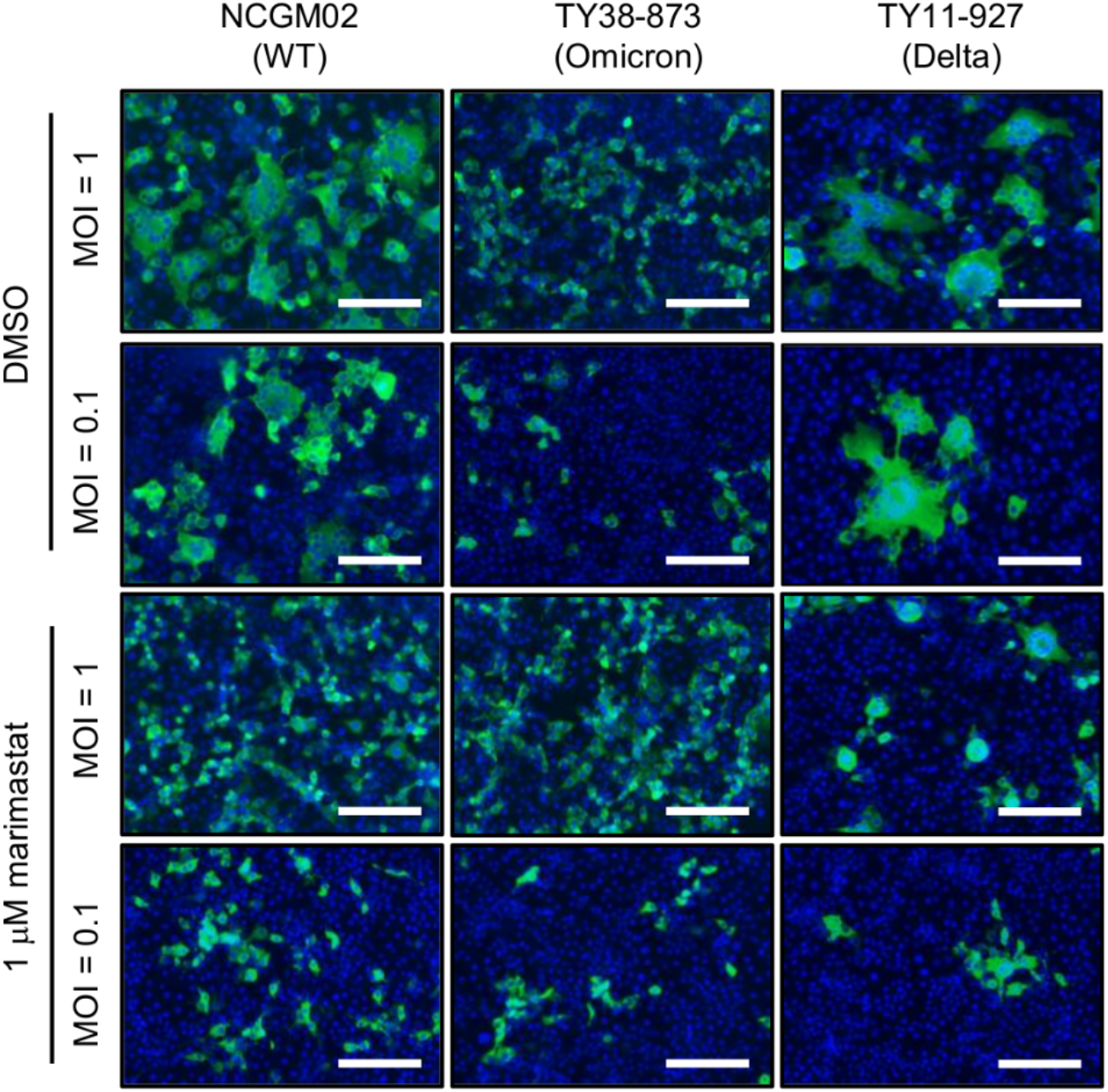
Effects of marimastat on syncytia formation of HEC50B cells induced by infection with WT, Omicron, and Delta virus. HEC50B cells were pretreated for 1 h with 1 μM marimastat and infected with the WT virus, Omicron, or Delta at an MOI of 0.1 or 1. After 24 h of infection, the cells were fixed and stained with anti-SARS-CoV-2 N antibody (green). Cell nuclei were stained with Hoechst 33342 (blue). White bar = 200 μm.

**SFig 4.**
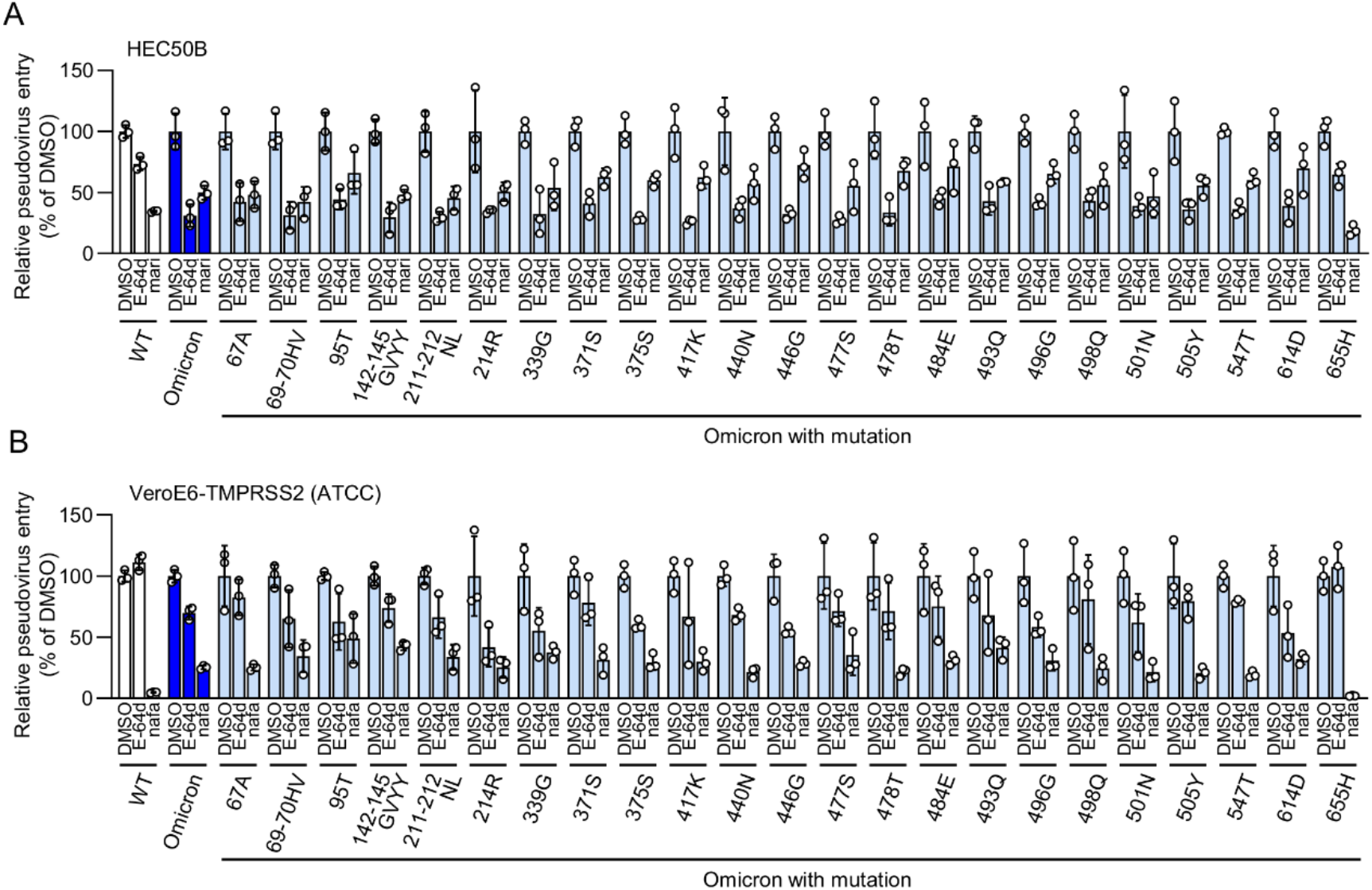
Effects of protease inhibitors on infection with VSV pseudoviruses bearing Omicron S proteins with each mutated amino acid residue in Omicron S1 substituted with the corresponding amino acid in WT S protein. Effects of protease inhibitors on infection with VSV pseudoviruses bearing Omicron S proteins with each mutated amino acid residue in the Omicron S1 substituted with the corresponding amino acid in the WT S protein. The relative pseudovirus entry was calculated by normalizing the luciferase activity for each condition to the luciferase activity of cells infected with pseudovirus in the presence of DMSO alone, which was set to 100%. Values are presented as mean ± SD (*n* = 3/group). *p > 0.05, **p > 0.01.

**SFig 5.**
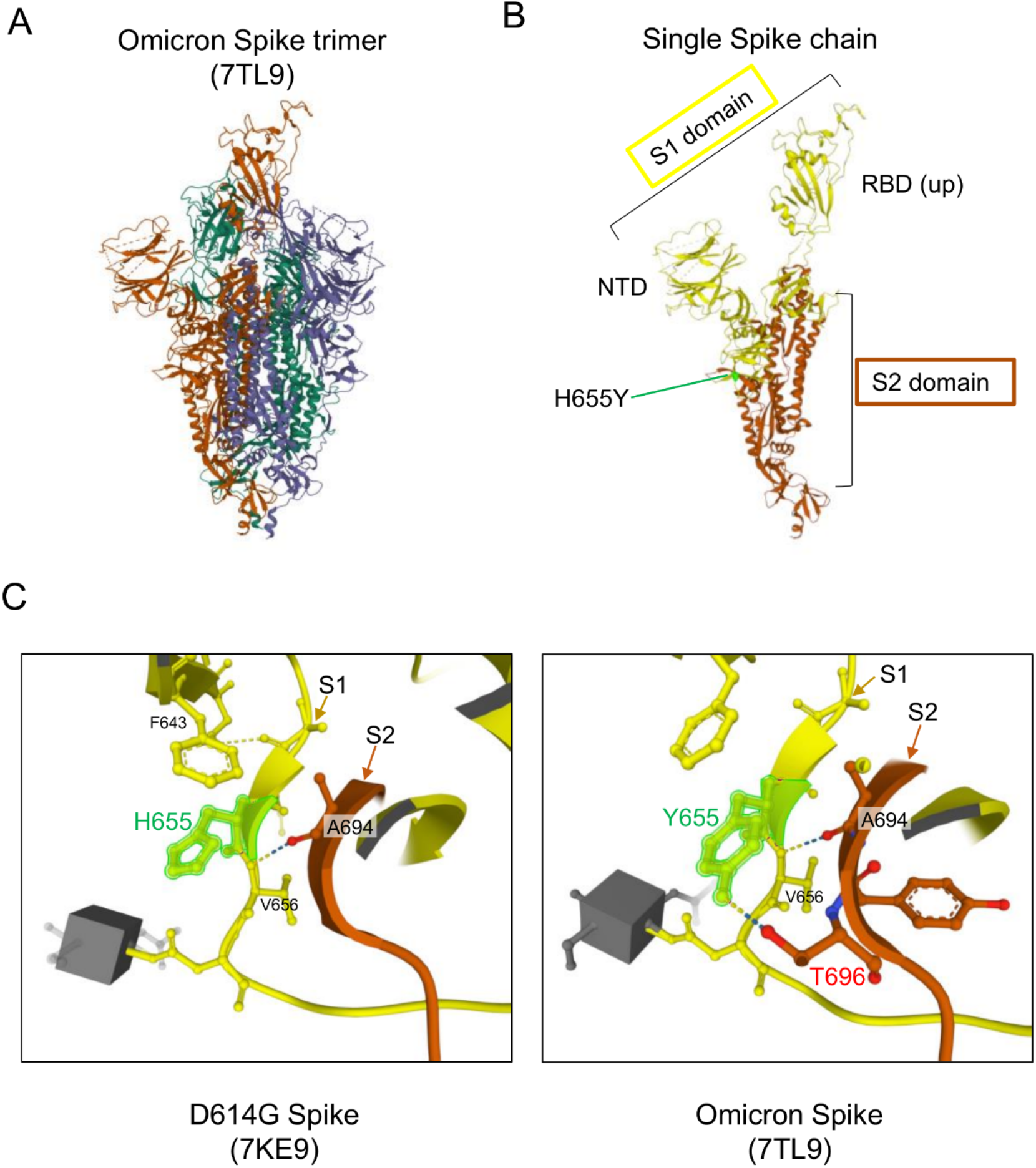
Prediction of additional hydrogen bond formation between tyrosine residue at position 655 (Y655) of Omicron S1 domain and threonine residue at position 696 (T696) in S2 domain. A. Three-dimensional structure of Omicron spike trimer (Protein Data Bank: 7TL9). B. Three-dimensional structure of a single-spike chain. S1 domain containing receptor binding domain (RBD), N-terminal domain (NTD), and S2 domain are indicated. The tyrosine residue at position 655 (Y655) was located in the binding region of the S1 and S2 domains. C. Structures of WT Spike (Protein data bank: 7KE9) and Omicron Spike (Protein data bank: 7TL9) around amino acid position 655 are shown. Both Spike proteins have hydrogen bonds between the two main chains (position 655-656 in S1 and position 694-695 in S2). Omicron Spike has an additional hydrogen bond between the two side chains (tyrosine residue at position 655 (Y655) in S1 and threonine residue at position 696 (T696) in S2).

## References

1. Halfmann PJ, Iida S, Iwatsuki-Horimoto K, Maemura T, Kiso M, Scheaffer SM, et al. SARS-CoV-2 Omicron virus causes attenuated disease in mice and hamsters. Nature. 2022. Epub 20220121. doi: 10.1038/s41586-022-04441-6. PubMed PMID: 35062015.

2. Suzuki R, Yamasoba D, Kimura I, Wang L, Kishimoto M, Ito J, et al. Attenuated fusogenicity and pathogenicity of SARS-CoV-2 Omicron variant. Nature. 2022. Epub 20220201. doi: 10.1038/s41586-022-04462-1. PubMed PMID: 35104835.

3. Shuai H, Chan JF, Hu B, Chai Y, Yuen TT, Yin F, et al. Attenuated replication and pathogenicity of SARS-CoV-2 B.1.1.529 Omicron. Nature. 2022. Epub 20220121. doi: 10.1038/s41586-022-04442-5. PubMed PMID: 35062016.

4. Tiecco G, Storti S, Degli Antoni M, Focà E, Castelli F, Quiros-Roldan E. Omicron Genetic and Clinical Peculiarities That May Overturn SARS-CoV-2 Pandemic: A Literature Review. Int J Mol Sci. 2022;23(4). Epub 20220211. doi: 10.3390/ijms23041987. PubMed PMID: 35216104; PubMed Central PMCID: PMCPMC8876558.

5. Garcia-Beltran WF, St Denis KJ, Hoelzemer A, Lam EC, Nitido AD, Sheehan ML, et al. mRNA-based COVID-19 vaccine boosters induce neutralizing immunity against SARS-CoV-2 Omicron variant. Cell. 2022;185(3):457-66.e4. Epub 20220106. doi: 10.1016/j.cell.2021.12.033. PubMed PMID: 34995482; PubMed Central PMCID: PMCPMC8733787.

6. Takashita E, Kinoshita N, Yamayoshi S, Sakai-Tagawa Y, Fujisaki S, Ito M, et al. Efficacy of Antibodies and Antiviral Drugs against Covid-19 Omicron Variant. N Engl J Med. 2022. Epub 20220126. doi: 10.1056/NEJMc2119407. PubMed PMID: 35081300; PubMed Central PMCID: PMCPMC8809508.

7. Hoffmann M, Krüger N, Schulz S, Cossmann A, Rocha C, Kempf A, et al. The Omicron variant is highly resistant against antibody-mediated neutralization: Implications for control of the COVID-19 pandemic. Cell. 2022;185(3):447-56.e11. Epub 20211224. doi: 10.1016/j.cell.2021.12.032. PubMed PMID: 35026151; PubMed Central PMCID: PMCPMC8702401.

8. Cameroni E, Bowen JE, Rosen LE, Saliba C, Zepeda SK, Culap K, et al. Broadly neutralizing antibodies overcome SARS-CoV-2 Omicron antigenic shift. Nature. 2022;602(7898):664–70. Epub 20211223. doi: 10.1038/s41586-021-04386-2. PubMed PMID: 35016195.

9. Dejnirattisai W, Huo J, Zhou D, Zahradník J, Supasa P, Liu C, et al. SARS-CoV-2 Omicron-B.1.1.529 leads to widespread escape from neutralizing antibody responses. Cell. 2022;185(3):467-84.e15. Epub 20220104. doi: 10.1016/j.cell.2021.12.046. PubMed PMID: 35081335; PubMed Central PMCID: PMCPMC8723827.

10. Zhao H, Lu L, Peng Z, Chen LL, Meng X, Zhang C, et al. SARS-CoV-2 Omicron variant shows less efficient replication and fusion activity when compared with Delta variant in TMPRSS2-expressed cells. Emerg Microbes Infect. 2022;11(1):277–83. doi: 10.1080/22221751.2021.2023329. PubMed PMID: 34951565; PubMed Central PMCID: PMCPMC8774049.

11. Hui KPY, Ho JCW, Cheung MC, Ng KC, Ching RHH, Lai KL, et al. SARS-CoV-2 Omicron variant replication in human bronchus and lung ex vivo. Nature. 2022. Epub 20220201. doi: 10.1038/s41586-022-04479-6. PubMed PMID: 35104836.

12. Meng B, Abdullahi A, Ferreira IATM, Goonawardane N, Saito A, Kimura I, et al. Altered TMPRSS2 usage by SARS-CoV-2 Omicron impacts tropism and fusogenicity. Nature. 2022. Epub 20220201. doi: 10.1038/s41586-022-04474-x. PubMed PMID: 35104837.

13. Gupta R. SARS-CoV-2 Omicron spike mediated immune escape and tropism shift. Res Sq. 2022. Epub 20220117. doi: 10.21203/rs.3.rs-1191837/v1. PubMed PMID: 35075452; PubMed Central PMCID: PMCPMC8786230.

14. Jackson CB, Farzan M, Chen B, Choe H. Mechanisms of SARS-CoV-2 entry into cells. Nat Rev Mol Cell Biol. 2021. Epub 20211005. doi: 10.1038/s41580-021-00418-x. PubMed PMID: 34611326; PubMed Central PMCID: PMCPMC8491763.

15. Hoffmann M, Kleine-Weber H, Pöhlmann S. A Multibasic Cleavage Site in the Spike Protein of SARS-CoV-2 Is Essential for Infection of Human Lung Cells. Mol Cell. 2020;78(4):779-84.e5. Epub 20200501. doi: 10.1016/j.molcel.2020.04.022. PubMed PMID: 32362314; PubMed Central PMCID: PMCPMC7194065.

16. Shang J, Wan Y, Luo C, Ye G, Geng Q, Auerbach A, et al. Cell entry mechanisms of SARS-CoV-2. Proc Natl Acad Sci U S A. 2020;117(21):11727–34. Epub 2020/05/06. doi: 10.1073/pnas.2003138117. PubMed PMID: 32376634.

17. Yan R, Zhang Y, Li Y, Xia L, Guo Y, Zhou Q. Structural basis for the recognition of SARS-CoV-2 by full-length human ACE2. Science. 2020;367(6485):1444–8. Epub 20200304. doi: 10.1126/science.abb2762. PubMed PMID: 32132184; PubMed Central PMCID: PMCPMC7164635.

18. Lan J, Ge J, Yu J, Shan S, Zhou H, Fan S, et al. Structure of the SARS-CoV-2 spike receptor-binding domain bound to the ACE2 receptor. Nature. 2020;581(7807):215–20. Epub 20200330. doi: 10.1038/s41586-020-2180-5. PubMed PMID: 32225176.

19. Hoffmann M, Kleine-Weber H, Schroeder S, Krüger N, Herrler T, Erichsen S, et al. SARS-CoV-2 Cell Entry Depends on ACE2 and TMPRSS2 and Is Blocked by a Clinically Proven Protease Inhibitor. Cell. 2020;181(2):271-80.e8. Epub 2020/03/05. doi: 10.1016/j.cell.2020.02.052. PubMed PMID: 32142651; PubMed Central PMCID: PMCPMC7102627.

20. Zhao MM, Yang WL, Yang FY, Zhang L, Huang WJ, Hou W, et al. Cathepsin L plays a key role in SARS-CoV-2 infection in humans and humanized mice and is a promising target for new drug development. Signal Transduct Target Ther. 2021;6(1):134. Epub 20210327. doi: 10.1038/s41392-021-00558-8. PubMed PMID: 33774649; PubMed Central PMCID: PMCPMC7997800.

21. Yamamoto M, Gohda J, Kobayashi A, Tomita K, Hirayama Y, Koshikawa N, et al. Metalloproteinase-dependent and TMPRSS2-independnt cell surface entry pathway of SARS-CoV-2 requires the furin-cleavage site and the S2 domain of spike protein. bioRxiv. 2021:2021.12.14.472513. doi: 10.1101/2021.12.14.472513.

22. Daniloski Z, Jordan TX, Wessels HH, Hoagland DA, Kasela S, Legut M, et al. Identification of Required Host Factors for SARS-CoV-2 Infection in Human Cells. Cell. 2021;184(1):92-105.e16. Epub 20201024. doi: 10.1016/j.cell.2020.10.030. PubMed PMID: 33147445; PubMed Central PMCID: PMCPMC7584921.

23. Carapito R, Li R, Helms J, Carapito C, Gujja S, Rolli V, et al. Identification of driver genes for critical forms of COVID-19 in a deeply phenotyped young patient cohort. Sci Transl Med. 2021:eabj7521. Epub 20211026. doi: 10.1126/scitranslmed.abj7521. PubMed PMID: 34698500.

24. Zhang Z, Zheng Y, Niu Z, Zhang B, Wang C, Yao X, et al. SARS-CoV-2 spike protein dictates syncytium-mediated lymphocyte elimination. Cell Death Differ. 2021;28(9):2765–77. Epub 20210420. doi: 10.1038/s41418-021-00782-3. PubMed PMID: 33879858; PubMed Central PMCID: PMCPMC8056997.

25. Bussani R, Schneider E, Zentilin L, Collesi C, Ali H, Braga L, et al. Persistence of viral RNA, pneumocyte syncytia and thrombosis are hallmarks of advanced COVID-19 pathology. EBioMedicine. 2020;61:103104. Epub 20201103. doi: 10.1016/j.ebiom.2020.103104. PubMed PMID: 33158808; PubMed Central PMCID: PMCPMC7677597.

26. Saito A, Irie T, Suzuki R, Maemura T, Nasser H, Uriu K, et al. Enhanced fusogenicity and pathogenicity of SARS-CoV-2 Delta P681R mutation. Nature. 2022;602(7896):300–6. Epub 20211125. doi: 10.1038/s41586-021-04266-9. PubMed PMID: 34823256; PubMed Central PMCID: PMCPMC8828475.

27. Rajah MM, Hubert M, Bishop E, Saunders N, Robinot R, Grzelak L, et al. SARS-CoV-2 Alpha, Beta, and Delta variants display enhanced Spike-mediated syncytia formation. EMBO J. 2021;40(24):e108944. Epub 20211025. doi: 10.15252/embj.2021108944. PubMed PMID: 34601723; PubMed Central PMCID: PMCPMC8646911.

28. Yamamoto M, Kiso M, Sakai-Tagawa Y, Iwatsuki-Horimoto K, Imai M, Takeda M, et al. The Anticoagulant Nafamostat Potently Inhibits SARS-CoV-2 S Protein-Mediated Fusion in a Cell Fusion Assay System and Viral Infection In Vitro in a Cell-Type-Dependent Manner. Viruses. 2020;12(6). Epub 2020/06/10. doi: 10.3390/v12060629. PubMed PMID: 32532094.

29. Yamamoto M, Matsuyama S, Li X, Takeda M, Kawaguchi Y, Inoue JI, et al. Identification of Nafamostat as a Potent Inhibitor of Middle East Respiratory Syndrome Coronavirus S Protein-Mediated Membrane Fusion Using the Split-Protein-Based Cell-Cell Fusion Assay. Antimicrob Agents Chemother. 2016;60(11):6532–9. Epub 2016/08/24. doi: 10.1128/aac.01043-16. PubMed PMID: 27550352; PubMed Central PMCID: PMCPMC5075056.

30. Johnson BA, Xie X, Bailey AL, Kalveram B, Lokugamage KG, Muruato A, et al. Loss of furin cleavage site attenuates SARS-CoV-2 pathogenesis. Nature. 2021;591(7849):293–9. Epub 20210125. doi: 10.1038/s41586-021-03237-4. PubMed PMID: 33494095; PubMed Central PMCID: PMCPMC8175039.

31. Chan JF, Zhang AJ, Yuan S, Poon VK, Chan CC, Lee AC, et al. Simulation of the Clinical and Pathological Manifestations of Coronavirus Disease 2019 (COVID-19) in a Golden Syrian Hamster Model: Implications for Disease Pathogenesis and Transmissibility. Clin Infect Dis. 2020;71(9):2428–46. doi: 10.1093/cid/ciaa325. PubMed PMID: 32215622; PubMed Central PMCID: PMCPMC7184405.

32. Baum A, Fulton BO, Wloga E, Copin R, Pascal KE, Russo V, et al. Antibody cocktail to SARS-CoV-2 spike protein prevents rapid mutational escape seen with individual antibodies. Science. 2020;369(6506):1014–8. Epub 20200615. doi: 10.1126/science.abd0831. PubMed PMID: 32540904; PubMed Central PMCID: PMCPMC7299283.

33. Dieterle ME, Haslwanter D, Bortz RH, Wirchnianski AS, Lasso G, Vergnolle O, et al. A Replication-Competent Vesicular Stomatitis Virus for Studies of SARS-CoV-2 Spike-Mediated Cell Entry and Its Inhibition. Cell Host Microbe. 2020;28(3):486-96.e6. Epub 20200703. doi: 10.1016/j.chom.2020.06.020. PubMed PMID: 32738193; PubMed Central PMCID: PMCPMC7332447.

34. Braun KM, Moreno GK, Halfmann PJ, Hodcroft EB, Baker DA, Boehm EC, et al. Transmission of SARS-CoV-2 in domestic cats imposes a narrow bottleneck. PLoS Pathog. 2021;17(2):e1009373. Epub 20210226. doi: 10.1371/journal.ppat.1009373. PubMed PMID: 33635912; PubMed Central PMCID: PMCPMC7946358.

35. Escalera A, Gonzalez-Reiche AS, Aslam S, Mena I, Pearl RL, Laporte M, et al. SARS-CoV-2 variants of concern have acquired mutations associated with an increased spike cleavage. bioRxiv. 2021:2021.08.05.455290. doi: 10.1101/2021.08.05.455290.

36. Bestle D, Heindl MR, Limburg H, Van Lam van T, Pilgram O, Moulton H, et al. TMPRSS2 and furin are both essential for proteolytic activation of SARS-CoV-2 in human airway cells. Life Sci Alliance. 2020;3(9). Epub 20200723. doi: 10.26508/lsa.202000786. PubMed PMID: 32703818; PubMed Central PMCID: PMCPMC7383062.

37. Yurkovetskiy L, Wang X, Pascal KE, Tomkins-Tinch C, Nyalile TP, Wang Y, et al. Structural and Functional Analysis of the D614G SARS-CoV-2 Spike Protein Variant. Cell. 2020;183(3):739-51.e8. Epub 20200915. doi: 10.1016/j.cell.2020.09.032. PubMed PMID: 32991842; PubMed Central PMCID: PMCPMC7492024.

38. Benton DJ, Wrobel AG, Xu P, Roustan C, Martin SR, Rosenthal PB, et al. Receptor binding and priming of the spike protein of SARS-CoV-2 for membrane fusion. Nature. 2020;588(7837):327–30. Epub 20200917. doi: 10.1038/s41586-020-2772-0. PubMed PMID: 32942285; PubMed Central PMCID: PMCPMC7116727.

39. Berger I, Schaffitzel C. The SARS-CoV-2 spike protein: balancing stability and infectivity. Cell Res. 2020;30(12):1059–60. doi: 10.1038/s41422-020-00430-4. PubMed PMID: 33139926; PubMed Central PMCID: PMCPMC7604330.

40. Gobeil SM, Henderson R, Stalls V, Janowska K, Huang X, May A, et al. Structural diversity of the SARS-CoV-2 Omicron spike. bioRxiv. 2022. Epub 20220126. doi: 10.1101/2022.01.25.477784. PubMed PMID: 35118469; PubMed Central PMCID: PMCPMC8811902.

41. Imai M, Iwatsuki-Horimoto K, Hatta M, Loeber S, Halfmann PJ, Nakajima N, et al. Syrian hamsters as a small animal model for SARS-CoV-2 infection and countermeasure development. Proc Natl Acad Sci U S A. 2020;117(28):16587–95. Epub 20200622. doi: 10.1073/pnas.2009799117. PubMed PMID: 32571934; PubMed Central PMCID: PMCPMC7368255.

42. Kandeel M, Yamamoto M, Tani H, Kobayashi A, Gohda J, Kawaguchi Y, et al. Discovery of New Fusion Inhibitor Peptides against SARS-CoV-2 by Targeting the Spike S2 Subunit. Biomol Ther (Seoul). 2021;29(3):282–9. doi: 10.4062/biomolther.2020.201. PubMed PMID: 33424013; PubMed Central PMCID: PMCPMC8094075.

